# Carriage of a single strain of non-toxigenic Corynebacterium diphtheriae biovar Belfanti (*Corynebacterium belfantii*) in four patients with cystic fibrosis

**DOI:** 10.1101/516880

**Authors:** Diane Pivot, Annlyse Fanton, Edgar Badell-Ocando, Marion Benouachkou, Karine Astruc, Frederic Huet, Lucie Amoureux, Catherine Neuwirth, Alexis Criscuolo, Serge Aho, Julie Toubiana, Sylvain Brisse

## Abstract

Cystic fibrosis (CF) patients are commonly colonized by bacterial pathogens, which can induce persistent lung inflammation and may contribute to clinical deterioration. Colonization of CF patients and cross-transmission by *Corynebacterium diphtheriae* has not been reported so far. The aim of this article was to investigate the possibility of a cross transmission of *C. diphtheriae* biovar Belfanti between four patients of a CF center. *C. diphtheriae* biovar belfanti (now formally called *C. belfantii*) isolates were collected from four patients in a single CF care center over a 6 years period and analyzed by microbiological methods and whole genome sequencing. Epidemiological links among patients were investigated. Ten isolates were collected from 4 patients. Whole genome sequencing of one isolate from each patient showed that a single strain was shared among them. In addition, one patient had the same strain on two consecutive samplings nine months apart. The strain was non-toxigenic and was susceptible to most antimicrobial agents. Ciprofloxacin resistance was observed in one patient. Transmission of the strain among patients was supported by the occurrence of same-day visits to the CF center. This study demonstrates colonization of CF patients by *C. diphtheriae* biovar Belfanti (*C. belfantii*) and shows persistence and transmission of a unique strain during at least six years in a single CF patient care center.

**Previous meeting presentations:** The information in this work was not previously presented in any meeting.

**Accession numbers:** The genomic sequence data generated in this work were submitted to the European Nucleotide Archive and are available from the International Nucleotide Sequence Database Collaboration (NCBI/ENA/DDBJ) databases under project accession number PRJEB28372 and run data accession numbers ERR2757916 to ERR2757921.

## INTRODUCTION

A large fraction of the mortality of cystic fibrosis (CF) patients is attributed to infections of the respiratory tract, which can be caused by multiple pathogens, and cross-transmission within CF centers themselves is an important healthcare related issue (1, 2)(3). The genus *Corynebacterium* includes a high number of pathogens, most of them being opportunistic (4). So far, only *C. pseudodiphtheriticum, C. propinquum and C. accolens* were reported from CF patients (5–7). *C. diphtheriae*, the most pathogenic *Corynebacterium* species that causes diphtheria, was not reported in pauci- or asymptomatic CF lung colonization to our knowledge. Typical diphtheria is caused by strains that produce the diphtheria toxin. Although the disease has almost disappeared in countries with high toxoid vaccine coverage, the pathogen still circulates in the human population (8–10). Further, non-toxigenic *C. diphtheriae* strains can be recovered from a variety of infections including respiratory tract infections, skin infections and bacteremia (11, 12). Three biovars of *C. diphtheriae* are distinguished by biochemical characteristics. Whereas biovars Mitis and Gravis can harbor the diphtheria toxin gene, biovar Belfanti isolates were only exceptionally described as toxigenic (13–15). Recently, *C. diphtheriae* biovar Belfanti isolates were recognized as representing a novel species called *C. belfantii* (16). The aim of this study was to investigate potential cross-transmission within a group of four patients with lower respiratory tract colonization by nontoxigenic *C. diphtheriae.* These patients were followed in a single CF center during a period of six years, and our genomic analyses showed that they were colonized by a single *C. diphtheriae* strain.

## METHODS

### Identification of cases

Cases were identified in the regional consultation CF Center of a university hospital between January 2011 and November 2016. During their visits at the center, patients are screened for the presence of opportunistic infectious agents and the evolution of antimicrobial resistance is monitored. The inclusion of cases was performed retrospectively based on at least one sample positive for *C. diphtheriae* upon microbiological screening from sputum or induced sputum. Clinical and laboratory data (sex, age at the time of diagnosis, pulmonary functionality, long-term or sequential antibiotic therapies, symptomatology at the time of diagnosis, respiratory co-infections) were collected for each patient.

### Epidemiological investigations

The CF center includes three wards (consultation, pulmonary function testing and chest physiotherapy) and an imaging department. The timeline of patients visits to the various wards of the CF center was investigated from their clinical records. Patients undergo chest physiotherapy either before or after their consultation with the physician, and a sputum specimen was systematically collected. Afterwards, they are directed to the pulmonary function testing ward.

To investigate infection control measures and detect possible factors that might have favored patient-to-patient transmission, all healthcare workers of the three wards who had worked during the study period were met and interviewed about implementation of standard precautions, material management and patient care organization. Besides, healthcare workers who were in charge of the included patients were screened based on a voluntary basis at the time of the study (June to October 2017) for *C. diphtheriae* colonization in the nasopharynx. For this purpose, swabs were plated onto blood agar medium, on which five fosfomycin disks (50 µg/disk) were then deposited. Colonies growing around the disks after 18-24 h at 37°C were sub-cultivated on Tinsdale medium agar and incubated at 37°C.

### Bacterial identification and characterization

The isolates were identified as *C. diphtheriae* at the local microbiology laboratory by MALDI-TOF (Bruker) since Sept. 2016, or using API Coryne (bioMerieux) until August 2016. Confirmatory analysis and *tox* gene detection were performed at the National Reference Center. The biovar of isolates was determined based on the combination of nitrate reductase (positive in Mitis and Gravis, negative in Belfanti) and glycogen fermentation (positive in Gravis only). Antimicrobial susceptibility was characterized by the disk diffusion method (Bio-Rad, Marnes-la-Coquette, France) and the minimum inhibitory concentration was determined by the E-test method (BioMérieux, Marcy l’Etoile, France). The sensitivity was interpreted using CA-SFM/EUCAST V.1.0 (mars 2017) criteria for *Corynebacterium* (http://www.sfm-microbiologie.org/UserFiles/files/casfm/CASFMV1_0_MARS_2017.pdf). Susceptibility was tested for the following antimicrobial agents: vancomycin, kanamycin, gentamycin, penicillin G, oxacillin, amoxicillin, imipenem, cefotaxime, clindamycin, azithromycin, spiramycin, clarithromycin, erythromycin, ciprofloxacin, moxifloxacin, cotrimoxazole, trimethoprim, sulfonamide, pristinamycin, rifampicin, tetracyclin and linezolid. Genomic sequencing was performed using a NextSeq-500 instrument (Illumina, San Diego, USA) with a 2 x 150 nt paired-end protocol and based on Nextera libraries. Contig sequences were assembled using SPAdes v3.9 (http://cab.spbu.ru/software/spades/). Multilocus sequence typing (MLST) was performed from genomic assemblies using the international nomenclature database webpage (https://pubmlst.org/cdiphtheriae/). Read mapping and calling of high quality single nucleotide polymorphisms (SNP) were performed as described (17). Identification to *C. belfantii* was performed by genomic comparison with the type strains of *C. belfantii* and *C. diphtheriae* based on the average nucleotide identity metric calculated with JspeciesWS (18) as in (16).

### Ethical statement

The work was conducted in accordance with local and national regulation, as well as the Helsinki Declaration, and was approved by the local Ethics committee (Committee for the Protection of Persons EST I, France).

## RESULTS

### Patients

From January 2011 to November 2016, four patients of the CF Center had positive screening respiratory samples for *C. diphtheriae*. This species was identified from these patients because of the presence of abundant, even though not numerically dominant, coryneform gram-positive colonies in their oropharyngeal microbiological flora. The timeline of patients visits to the CF center and detection of *C. diphtheriae* is represented in **Figure 1**. The patient characteristics and medical records are summarized in **Table 1**. Only patients 2 and 4 presented with respiratory exacerbation at the time of *C. diphtheriae* detection and received antibiotics, but they did not require hospitalization. None of the four patients had a dermatological disease or chronic wound. Patient 2 was positive for *C. diphtheriae* during at least 15 months.

**Table 1.** Summary of the medical records of the four patients with positive *C. diphtheriae* isolates in expectoration samples

| Patient | Age/<br>Gen<br>der | Baseline lung<br>function | Symptoms at the time<br>of positivity | Long-term or recurrent<br>antibiotic therapy | Date of<br>the first<br>positive<br>sample | Sampling | Other bacteria<br>(co-infection) | Treatment<br>following<br>positivity | Current<br>status |
| --- | --- | --- | --- | --- | --- | --- | --- | --- | --- |
| 1 | 27/F | Non-invasive<br>ventilation, oxygen<br>requirement<br>FEV 30% | Chronic respiratory<br>failure | Amoxicillin,<br>cotrimoxazole for<br>exacerbations | Feb 2011 | Sputum | *S. maltophilia*<br>*H. influenzae*<br>*E. coli* | No treatment | Deceased in<br>2013 |
| 2 | 25/<br>M | FEV 106% | Asymptomatic in Apr<br>2015,<br>Exacerbation in Jul 2015 | Inhaled tobramycin for<br>one month on Jan 2014;<br>Ofloxacin, fusidic acid | Apr 2015 | Sputum | *S. aureus* | Fusidic acid,<br>ofloxacin | Alive |
| 3 | 39/<br>M | FEV 69% | Asymptomatic | amoxicillin/clavulanic<br>acid, ofloxacin, amikacin<br>for exacerbation | Jan 2016 | Sputum | *S. aureus*<br>*A. baumannii*<br>*S. maltophilia* | No treatment | Alive |
| 4 | 23/<br>M | FEV 75% | Exacerbation in Nov 2016 | Piperacillin/tazobactam,<br>teicoplanin, imipenem,<br>cotrimoxazole for<br>exacerbation in 2015 and<br>2016 | Nov 2016 | Sputum | *S. aureus*<br>*Achromobacter*<br>*xylosoxidans* | Amoxicillin/<br>clavulanic acid,<br>cotrimoxazole | Alive |

FEV: force expiration volume

**Figure 1.**
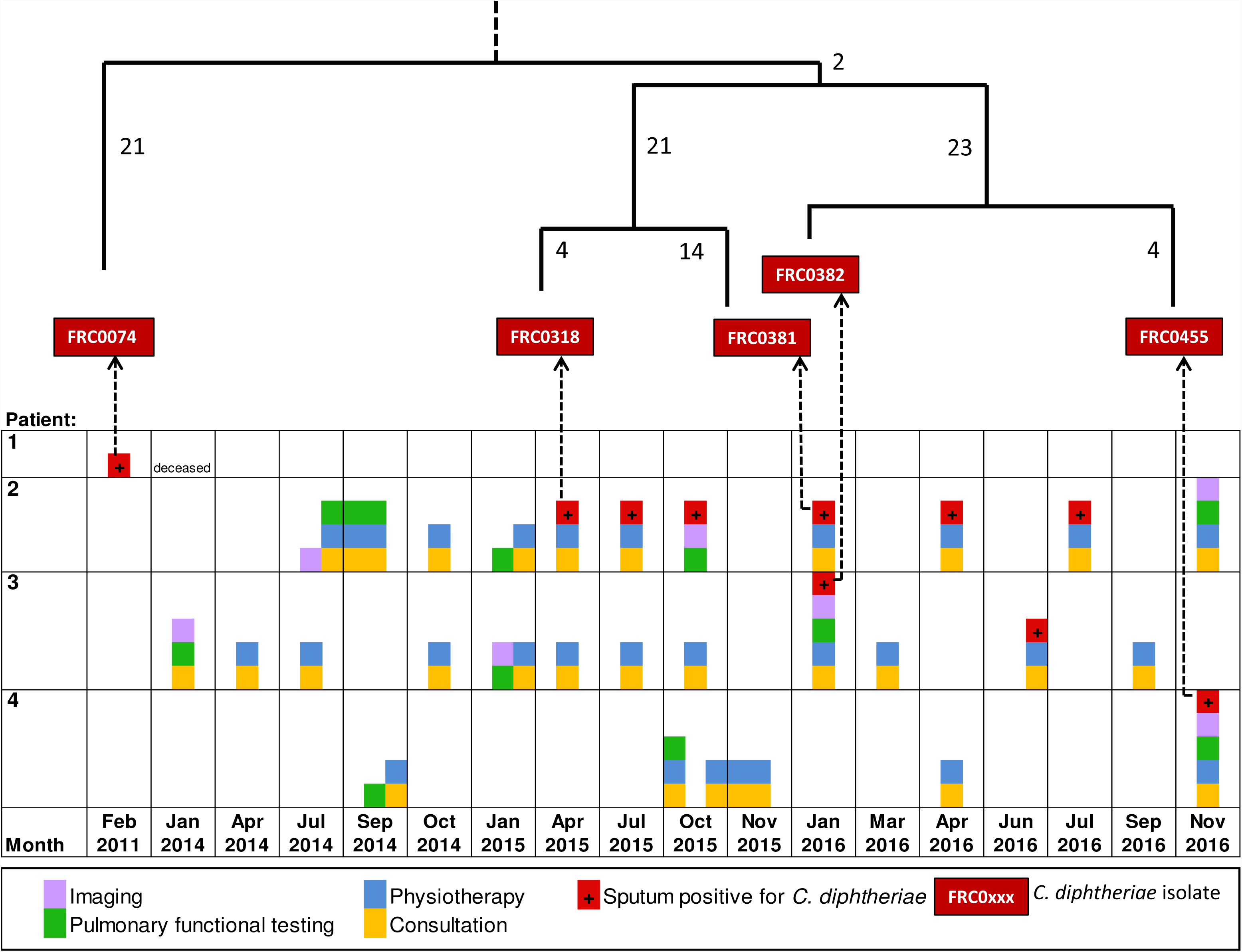
Timeline of events and *C. diphtheriae* detection in four cystic fibrosis patients. Months correspond to different columns in the grid; inside each month, days (not shown) are distinguished as separate columns. The upper tree recapitulates the phylogenetic relationships among the isolates, which are indicated in red boxes, linked with dotted arrows to their respective patient and isolation time. The numbers of single nucleotide polymorphisms (SNP) per genome inferred to have occurred are given for each branch of the tree.

### Phenotypic and molecular identification of the isolates

One isolate from patients 1 (February 2011), 3 (January 2016) and 4 (November 2016), and 2 isolates from patient 2 collected 9 months apart (April 2015, January 2016), were stored and available for analysis (**Figure 1**). The five isolates were identified as *C. diphtheriae.* None of the isolates was toxigenic and all were of biovar Belfanti. MLST showed that the five isolates belonged to the same sequence type, ST208. Whole-genome sequence variation among the 5 isolates revealed only 62 SNPs among them. The largest SNP distance between two isolates was 58 SNPs, observed between isolates FRC0074 and FRC0381. Hence, the five isolates were very closely related, showing that they belong to a single strain. Phylogenetic analysis based on SNPs (**Figure S1**) revealed three subtypes, comprising respectively (1) the isolate from patient 1 (FRC0074); (2) One isolate from patient 2 (isolate FRC0318) and the isolate from patient 3 (FRC0381); and (3) The other isolate (FRC0382) from patient 2 and the isolate from patient 4 (FRC0455). Within subtypes, only eighteen SNPs separated FRC0318 and FRC0381, and 4 SNPs separated FRC0382 and FRC0455. Following the recent description of *C. belfantii* (16), the five isolates were re-identified based on their genomic sequence. Their average nucleotide identity with *C. belfantii* type strain (FRC0043^T^) was 99.47%, whereas it was only 94.89% with NCTC11397^T^, the type strain of *C. diphtheriae*. Therefore, the five isolates belong to the novel species *C. belfantii*.

Antimicrobial susceptibility profiles of the five isolates showed that they were susceptible to all antimicrobial agents, with one remarkable exception: isolate FRC0074 from patient 1 was non-susceptible to ciprofloxacin. Genomic sequence inspection showed that this isolate had a unique mutation, A to G at position 277 of the *gyrA* gene, coding for subunit A of gyrase, the target of ciprofloxacin. This SNP corresponds to a deduced amino-acid change of D to N at protein position 91, which is located within the quinolone-resistance determining region of the gyrase of *Corynebacteria*.

### Investigations of possible strain transmission risks

Patient 1 had no recorded contact opportunity with the three other patients at the clinic. In contrast, patients 2 and 3 visited the CF center the same day on eight occasions for consultations, physiotherapy or for pulmonary function tests between January 2014 and November 2016 (**Figure 1**). Further, patient 4 had a consultation and physiotherapy session 30 minutes after patient 2 in April 2016 and then in November 2016, on the day when he had a positive expectoration sample for *C. diphtheriae*. In addition, patient 4 had a pulmonary function test on the same day as patient 2 in September 2014 and November 2016 (**Figure 1**). Therefore, several opportunities for cross-transmission within the CF center were identified among patients 2, 3 and 4, even though social interaction between CF patients was not identified.

Inspections of hygiene measures was retrospectively conducted by the infection control team between June 2016 and June 2017. Local infection control protocols and recommendations regarding healthcare staff hygiene (mostly masks, disposable mouthpieces and filters, specific equipment and hand hygiene) and rooms and equipment disinfection were correctly observed. However, it was noted that the salbutamol inhalation chamber was disinfected with low-level disinfectant instead of a mid-level disinfectant. Furthermore, after their physiotherapy session, patients did not always wear masks while undergoing pulmonary function testing.

Ten healthcare workers of the CF center who were regularly in contact with the four patients were retrospectively screened for throat colonization by *C. diphtheriae*. No *Corynebacterium* was isolated in any of the samples from the screened workers.

## DISCUSSION

We report four cases of CF patients colonization by non-toxigenic *C. diphtheriae* biovar Belfanti (now formally called *C. belfantii*). To our knowledge, *C. diphtheriae* was never described from CF patients. Although non-toxigenic isolates can cause a variety of infections, including bacteremia (11), two patients were not symptomatic, whereas the two other had lung exacerbation at the time of first *C. diphtheriae* detection. Other opportunist pathogens of the *Corynebacterium* genus might be involved in CF lung exacerbations (5, 19). In recent years, *C. diphtheriae* became easier to identify by the use of MALDI-TOF. However, *C. diphtheriae* may still be difficult to detect in the CF lungs, given the polymicrobial colonization occurring in most samples, as was observed during this study (**Table 1**). It is therefore not unlikely that additional cases of colonization might have gone undetected.

Multiple CF patients are typically followed in a given CF center, which creates opportunities for bacterial transmission among patients despite the enforcement of strong infection control measures. A strong suspicion of cross-transmission of *C. diphtheriae* between four patients in our clinic arose given the repeated observation of patients colonized by *C. diphtheriae*. Microbiological investigations fully supported the hypothesis of a single strain. MLST showed that the five isolates belonged to the same sequence type. The MLST genotype of the isolates, ST208, was not reported previously in the *C. diphtheriae* MLST database, suggesting that it is not common. Whole genome sequencing defines the genetic relatedness among *C. diphtheriae* isolates with high precision (20–22). This approach demonstrated that the five isolates belong to the same strain and provided strong support to the hypothesis of cross transmission among patients and/or contamination from a common source. Besides, the data showed that the strain persisted within patient 2 for at least 9 months. Unfortunately, the additional isolates detected from patient 2 and 3 were not stored.

The SNP variation uncovered by the genomic analysis reflects evolution of the strain since the last common ancestor of the five isolates. Three subtypes were distinguished, two of which comprised isolates from two patients. Subtypes shared by patients may reflect direct transmission between them. Epidemiological investigations support this possibility in one case, as patients 2 and 3 had simultaneous visits to the CF center on several occasions. However, knowledge on subtype diversity within patients would be required to infer transmission chains with confidence (23). In this study, only one isolate was kept and characterized from each sample. Therefore, one cannot exclude that subtypes coexisted within single patients, which would lead to the possibility of other transmission patterns than those suggested by the phylogeny.

Biovar Belfanti of *C. diphtheriae* (*C. belfantii*) is commonly isolated from the respiratory tract, generally from the nose or throat and often in association with ozaena (24). In contrast, its isolation from skin infections is extremely rare. Therefore, skin wounds of patients or the personnel is an unlikely reservoir. Transmission by direct respiratory contamination between patients appears as the most likely transmission route. Transmission of *C. striatum* in an intensive care unit and silent transmission of *C. pseudodiphtheriticum* among CF patients were previously reported (5, 25).

Evolution of antimicrobial resistance occurs frequently in bacterial isolates that colonize CF lungs (26). Our results showed ciprofloxacin resistance in one isolate, whereas the others were susceptible. As no prescription of quinolone antimicrobials was recorded for this patient, one possibility is that the strain had evolved resistance to ciprofloxacin in another ciprofloxacin-treated patient, in which the strain was not detected.

The main limitation of this study is that the infection control investigation was performed retrospectively and that no detailed pattern of transmission could therefore be ascertained. Healthcare workers or the materials used for patient care may have played the role of vector of *C. diphtheriae* between the patients (27), even though the retrospective screening did not reveal a potential carrier or source of infection. Further, hidden patient-to-patient cross transmission cannot be excluded, as some patients may have been undetected for *C. diphtheriae* carriage. Further studies are needed to better define carriage of *C. diphtheriae* by CF patients and to investigate the possible role of patients, healthcare workers or environmental sources in cross transmission. In addition, the clinical significance of non-toxigenic *C. diphtheriae* will need to be determined in order to define strategies of treatment, prevention and control of contamination of CF patients by this bacterial pathogen.

## Acknowledgements

We acknowledge the help of Melody Dazas and Annick Carmi-Leroy for the microbiological characterization of the isolates. We thank the “Plateforme de Microbiologie Mutualisée” from Institut Pasteur for genomic sequencing.

Abbreviations

CF: cystic fibrosis
MLST: multilocus sequence typing
MALDI-TOF: Matrix-assisted laser desorption/ionization time-of-flight

